# Accurate Microbiome Sequencing with Synthetic Long Read Sequencing

**DOI:** 10.1101/2020.10.02.324038

**Authors:** Nico Chung, Marc W. Van Goethem, Melanie A. Preston, Filip Lhota, Leona Cerna, Ferran Garcia-Pichel, Vanessa Fernandes, Ana Giraldo-Silva, Hee Shin Kim, Evan Hurowitz, Michael Balamotis, Indira Wu, Tuval Ben-Yehezkel

**Affiliations:** Welgene Biotech, Taipei, Taiwan; Lawrence Berkeley National Laboratory, Berkeley, CA, USA; Promega Corporation, Madison, WI, USA; Gennet, Centre for Medical Genetics and Reproductive Medicine, Prague, Czech Republic; Center for Fundamental and Applied Microbiomics, Arizona State University, AZ, USA; Loop Genomics, San Jose, CA, USA

## Abstract

The microbiome plays a central role in biochemical cycling and nutrient turnover of most ecosystems. Because it can comprise myriad microbial prokaryotes, eukaryotes and viruses, microbiome characterization requires high-throughput sequencing to attain an accurate identification and quantification of such co-existing microbial populations. Short-read next-generation-sequencing (srNGS) revolutionized the study of microbiomes and remains the most widely used approach, yet read lengths spanning only a few of the nine hypervariable regions of the 16S rRNA gene limit phylogenetic resolution leading to misclassification or failure to classify in a high percentage of cases. Here we evaluate a synthetic long-read (SLR) NGS approach for full-length 16S rRNA gene sequencing that is high-throughput, highly accurate and low-cost. The sequencing approach is amenable to highly multiplexed sequencing and provides microbiome sequence data that surpasses existing short and long-read modalities in terms of accuracy and phylogenetic resolution. We validated this commercially-available technology, termed LoopSeq, by characterizing the microbial composition of well-established mock microbiome communities and diverse real-world samples. SLR sequencing revealed differences in aquatic community complexity associated with environmental gradients, resolved species-level community composition of uterine lavage from subjects with histories of misconception and accurately detected strain differences, multiple copies of the 16S rRNA in a single strain’s genome, as well as low-level contamination in soil cyanobacterial cultures. This approach has implications for widespread adoption of high-resolution, accurate long-read microbiome sequencing as it is generated on popular short read sequencing platforms without the need for additional infrastructure.

## Introduction

Next-generation sequencing (NGS) has revolutionized microbiome research in the 21st century (Berg *et al*., 2020). Considerable recent research has leveraged its power, which, coupled with the remarkable decrease in sequencing costs over the past two decades, has led to an explosion in microbiome research, making it possible to access rare microbial community members (so-called ‘microbial dark matter’) (Turner *et al*., 2013, Lloyd-Price *et al*., 2016). As the fidelity and accuracy of these technologies continue to increase, so too do the number of microbiome-related hypotheses that become testable using NGS methodologies.

Characterizing the bacterial (and to a broader extent, the prokaryotic) component of the microbiome routinely relies on 16S ribosomal RNA (rRNA) gene sequencing. The 16S rRNA gene sequence, which contains nine hypervariable regions (V1-V9) in an ∼1,550 bp sequence, is used as a classifier to identify distinct phylogenies and document their relative abundances within mixed, complex populations, offering a window into community composition, contributing to our understanding of microbes and their relationship to human health and the environment, typically through high-throughput, cost-effective short-read NGS (srNGS)

However, srNGS sequencing suffers from a few notable shortcomings. Critically, even the longest paired-end reads generated by srNGS typically only sequence ∼500 bp fragments after merging reads (Martijn *et al*., 2019), which only cover a few (e.g. V3-V4) of the 16S rRNA’s nine hypervariable regions (or some 16-33% of its sequence) that allow to differentiate sequence variants for taxonomic classification. And yet, accurate classification of 16S rRNA gene sequences is advantageous for evaluating the composition of microbiome samples, yet short-read coverage of the 16S hypervariable regions typically only enables confident sequence classification at the genus or sometimes species leve<u>l</u> (Johnson *et al*., 2019). This often impedes interpretation of closely-related yet functionally distinct species. Moreover, the inability to resolve 16S rRNA gene copy numbers (GCNs) from mixed microbial populations biases sequence variant counts towards members that contain more GCNs per genome (Louca *et al*., 2018).

An alternative approach to overcome some of these limitations could be long-read sequencing. Platforms offered by Pacific Biosciences® (i.e. Single Molecule Real-Time sequencing, and Oxford Nanopore (such as the MinION) provide sequence data that encompasses regions V1 - V9 and dramatically improve taxonomic classification to the strain leve<u>l</u> (Benítez-Páez *et al*., 2016, Callahan *et al*., 2019). However, long-read technologies are not without shortcomings. Notably, long-read raw sequences harbor very high error rates (up to 15% of nucleotides per sequence) and have a lower throughput than short-read sequencing which can limit utility in large-scale studies. Furthermore, high costs impede their wider application (Kuleshov *et al*., 2016).

To circumvent these challenges, here we perform a technical evaluation of LoopSeq 16S sequencing, a synthetic long-read (SLR) technology that builds on previous work in the SLR domain (Hong *et al*., 2014, Stapleton *et al*., 2016) and addresses the shortcomings of short-read 16S rRNA gene sequencing for microbiome characterization. SLR sequencing leverages the high-throughput, base call accuracy and the low cost of short-read sequencing to generate long-read, high fidelity sequence data that capture all nine hypervariable regions in a single molecule readout, covering the full-length 16S ribosomal RNA gene. We evaluated LoopSeq 16S sequencing using a two-pronged approach. First, we validated the method’s reliability (error rates, composition bias, classification accuracy) using three commercially available microbiome mock communities with well-characterized bacterial compositions produced by two independent manufacturers. Second, we tested the method’s utility and performance using multiple samples from highly diverse natural environments including aquatic (lake, pond, and aquarium), uterine lavage microbiomes, and non-axenic cyanobacterial cultures isolated from biological soil crust communities (biocrusts). We then demonstrate how LoopSeq 16S sequencing can overcome some of the limitations associated with the use of traditional short read NGS approaches.

## Materials & Methods

### Mock community from Zymo Research

The ZymoBIOMICS^™^ Microbial Community Standard D6306 (termed ‘Zymo’ throughout) is commercially available from Zymo Research (Zymo Research, Orange, CA, USA) and comprises eight bacterial species and two fungal species. The fungal component of the standard was not sequenced in this study as our analysis is focused exclusively on sequencing the 16S rRNA gene.

### Mock communities from ATCC

Two mock community samples representing the human gut (termed ‘ATCC-gut’ throughout; MSA-1006) and the human oral cavity (termed ‘ATCC-oral’ throughout; MSA-1004) were used for microbiome analysis. Reference sequences were retrieved from American Type Culture Collection (ATCC); ATCC Genome Portal (https://genomes.atcc.org/). For any strains not in this database, we used the strain name to find identical or similar strain references in other genome repositories (i.e. NCBI RefSeq CG database).

### Environmental samples

#### Water samples

Four water samples were collected from several sources in Madison, WI, USA. Specifically, samples were collected from an aquarium (“Aquarium”), a pond (“FeynmanPond”, GPS coordinates 43°00’19.7” N, 89°25’10.8” W), a barrel collecting roof rain runoff (“RainBarrel”) and a garden pond (“RivaPond”, GPS coordinates 43°01’46.8” N, 89°29’01.6” W). The water samples were filtered through a 0.2 μm pore size polycarbonate disc filter (Whatman® Nuclepore™ Track-Etched Polycarbonate Membrane, Sigma) to collect microbial biomass. Filters were transferred to 2 mL microfuge tubes for further processing using the Maxwell® RSC PureFood GMO and Authentication Kit (Promega Corporation) with a modified version of the kit protocol. Following the addition of 1 mL of CTAB Buffer, samples were incubated at 95°C for 5 minutes, cooled for 2 minutes at room temperature, and vortexed for 1 minute. Samples were treated with RNase A and Proteinase K at 70°C for 10 minutes. DNA from 300µl of the resulting lysates was purified with the Maxwell® RSC Instrument according to manufacturer’s kit instructions. DNA eluates were cleaned up using Zymo Research OneStep PCR Inhibitor Removal Kit, according to manufacturer’s instructions.

Cleanup steps during LoopSeq library preparation were performed with ProNex® Size-Selective Purification System (Promega Corporation), using manufacturer’s binding and washing instructions and the elution volume specified in the LoopSeq kit. Specifically, the volumes of ProNex® Chemistry used for each cleanup step were as follows: 55.9 µl (1.075x) for post-barcoding cleanup, 132 µl (1.1x), 137.8 µl (1.3x) for post-activation cleanup, 135 µl (1.35x) for post-ligation cleanup, and 55 µl (1.1x) for post-indexing cleanup. ProNex® NGS Library Quant Kit (Promega Corporation) was used for library quantification prior to sequencing on a MiSeq® instrument. The library of pooled samples was denatured and diluted to 15 pM. PhiX was denatured and diluted to 20 pM and added to the library at 5% of the total diluted library volume, according to Illumina Document #1000000061014 v00. DNA samples were stored at -20°C until further processed for sequencing with LoopSeq kit.

#### Uterine lavage *samples*

Human-derived uterine lavage samples were prepared from two patients with complications related to conception. Patient 1: aged 48, secondary infertility (2x spontaneous abortion, 1x induced abortion for interstitial pregnancy, multiple unsuccessful attempts of assisted reproduction (6x embryo transfer, multiple intrauterine inseminations). Patient 2: aged 35, long-term unsuccessful effort for natural conception. To examine unique microbial communities from human-derived sources, the uterine lavage samples were prepared using the ZymoBIOMICS™ DNA Microprep Kit from Zymo Research (D4301). Notably, we included a mechanical lysis bead beating procedure to release genetic material from microbes that produce endospores (such as *Firmicutes*) that are recalcitrant to chemical lysis alone.

#### Cyanobacterial enrichment cultures from biological soil crust communities

Enrichment cultures for soil cyanobacteria were obtained and maintained in the lab according to Giraldo-Silva *et al*. (2019). Genomic DNA was extracted using the DNeasy® PowerSoil® Kit (Qiagen), reference number 12888-100, following manufacturer’s instructions from two non-axenic cyanobacterial enrichments from soils in the Great Basin Desert (Giraldo-Silva *et al*., 2019); HSN023 -Sample7 [well G1]) and from the Sonoran Desert (Fernandes *et al*. in prep; CYAN3 - Sample-16 [well H2]). Final eluted genomic DNA was stored at -20 °C until further process using the LoopSeq 16S long-read kit.

#### LoopSeq Library Sequencing

All mock community and environmental samples were prepared with LoopSeq kits and sequenced on the Illumina NextSeq 500 platform run in 150 bp paired-end mode. For each LoopSeq 16S kit, which can process up to 24 samples simultaneously and capturing ∼12,000 16S long-read per sample (∼300k 16S molecules from a complete kit run), 100-150M PE reads (50-75M clusters passing filter) were dedicated for each sequencing run, yielding ∼20 Gb of data. The Zymo-V3V4 and uterine lavage libraries were sequenced using the Illumina MiSeq platform run in 300 bp paired-end mode, yielding ∼15 Gb of data.

#### Library preparation of LoopSeq (V1-V9) and short-read (V3V4) libraries

Sequencing libraries were made from the Zymo mock community using the LoopSeq 16S long-read kit (referred to throughout as ‘Zymo-Loop’). This preparation was also followed for the ATCC-gut and ATCC-oral communities as well as the uterine lavage and cyanobacterial enrichment samples. For the water samples, libraries were made using the LoopSeq 16S & 18S long-read kit, which captures both prokaryotic and eukaryotic rDNA long-read targets, but only the assembled 16S contigs from the final sequencing output were analyzed here.

To compare LoopSeq SLR technology against Illumina-based short-read sequencing for classifying a bacterial population, a sequencing library that captured the V3V4 hypervariable region of the 16S rRNA gene was made from the Zymo mock community (named ‘Zymo-V3V4’). To generate this comparison library, primers that flanked the V3 and V4 16S gene region were used to directly PCR amplify from genomes in the Zymo mock community (primer sequence available upon request). The following PCR protocol was used to generate V3V4 amplicons: 95°C for 5 mins then 98°C for 20 s, 58°C for 20s, and 72°C for 30s for 25 cycles, followed by clean up with SPRI (0.8x) and elution in 50 μl of 10 mM Tris buffer, pH 8.0. Following elution, amplicons underwent a PCR-based addition of Illumina indexing adapters (P5-R1 and P7-index-R2; sequence available upon request). The following PCR protocol was used for indexing: 95°C for 3 minutes then (98°C for 30s, 58°C for 30s, 72°C for 30s) for 8 cycles and ending with a 72°C step for 5 minutes, followed by clean up with SPRI (1.2x) and final elution in 25 μl of 10mM Tris buffer, pH 8.0. For all PCR reactions, a KAPA 2x HiFi HotStart ReadyMix (Roche, 89125-040) was used. Detailed description of the materials and methods and procedures used to prepare the LoopSeq libraries are available at: https://bit.ly/3bQMAzt.

#### Bioinformatic analyses

For the Zymo-Loop data, full-length 16S contigs were filtered by matching to gene-specific forward (27F: AGAGTTTGATCMTGGCTCAG) and reverse (1492R: TACCTTGTTACGACTT) primer sequences using the DADA2 v1.14.0 package in R (Callahan *et al*., 2016). Zymo-Loop was mapped to the given mock community references using Bowtie2 v2.2.9 (Langmead & Salzberg, 2012) in end-to-end mode on the very-sensitive setting.

To determine the accuracy of LoopSeq SLR contigs, we examined the intragenomic copy ratios for the ten 16S rRNA genes present in the *B. subtilis* genome (with 5 non-redundant copies: #1, #4, #8, #9, #10 are identical; #2, #3 are identical; #5, #6, #7 are individually unique). Mapped contigs were extracted and matched to a sequence database containing the different copies of the *B. subtilis* 16S rRNA genes. The error profile was assessed with Alfred v0.1.17 (Rausch *et al*., 2019).

The Zymo-V3V4 short-reads were filtered for the PhiX spike-in sequence using Bowtie2 v2.2.9 (Langmead & Salzberg, 2012), quality filtered and trimmed using Trimmomatic v0.36 (Bolger *et al*., 2014). Paired reads were merged with FLASH v1.2.11 (Magoč & Salzberg, 2011) primers trimmed using Cutadapt (Martin, 2011), and filtered by length (min = 440 bp, max = 450 bp) using DADA2 v1.14.0. These processed fragments were then mapped to the Zymo whole genome references to determine relative species abundances against the known reference composition. The reads that mapped to *B. subtilis* were extracted and re-mapped to non-redundant *B. subtilis* 16S rRNA gene copy references using Bowtie2. The error profile was assessed using the sequence alignment QC program Alfred (Rausch *et al*., 2019).

The eight bacterial species in the Zymo mock community are sufficiently phylogenetically distant such that reads and contigs are unlikely to be erroneously mapped to the wrong species given their length and accuracy. However, it is often not the case that the species within a sample are known *a priori* while reference sequences for those species may be unavailable. It is more common that samples with unknown taxonomic composition must be compared to a more general database. To determine how well 16S rRNA sequences can be used to accurately identify the correct species using a general database, full-length Zymo-Loop contigs were mapped using Bowtie2 v2.2.9 to all bacterial genomes in the NCBI RefSeq CG (complete genomes) database (obtained October 2019). The database contains genomes of the same species as those in the Zymo sample although it is unclear if the strains are the same since this information is lacking. Importantly, there are many genomes of closely related species (i.e. from the same genus) that could potentially confound a direct mapping approach to species identification.

Whereas the species in the Zymo samples are all represented in NCBI RefSeq bacterial database, it is more likely the case that real samples will contain novel species. As such, it may be more pertinent to use *k*-mer based lowest common ancestor (LCA) classification methods. Novel species not represented in the database may nonetheless be classified with some degree of confidence to the genus taxonomic level. However, commonly used *k*-mer based LCA methods such as mothur (Schloss *et al*., 2009) or QIIME (Caporaso *et al*., 2010) use curated 16S rRNA gene databases such as SILVA (Pruesse *et al*., 2007), Greengenes (DeSantis *et al*., 2006), or the Ribosomal Database Project (RDP; (Cole *et al*., 2005) that typically do not contain all 16S rRNA gene copies from represented species which may lead to skewed estimates of taxonomy.

Assembled long contigs from the two ATCC mock communities were filtered for full-length 16S rRNA genes and mapped using Bowtie2 to exact strain references. The Madison water samples were filtered for full-length 16S rRNA genes and classified using Kraken 2 and Bracken with the standard database. Kraken 2 (Wood *et al*., 2019) is a *k*-mer based LCA method that can use whole genome references, thus allowing all 16S rRNA gene copies to be represented. The standard Kraken 2 database contains (accessed October 2019), the entire NCBI RefSeq complete genomes for bacteria, archaea, virus, and humans. Since Kraken 2 is an LCA method, it may assign sequences to any taxonomic level. Therefore, the results cannot necessarily provide overall species ratios with confidence. To evaluate the accuracy of mapping LoopSeq SLR contigs to expected species ratios, a companion program to Kraken 2, Bracken (Lu *et al*., 2017), was used to redistribute contigs to the appropriate species level.

The water samples, and uterine lavage full-length 16S rRNA gene sequences were classified with Kraken 2 and Bracken. The two biocrust cyanobacterial enrichments were sequenced on LoopSeq 16S architecture and processed as above. Here we consolidated all full-length ASVs to represent strain-level taxonomies at 100% nucleotide identity. Resulting feature tables, with full length 16S rRNA genes, were used to assign taxonomy to each of the 16S rRNA sequences obtained for each of the used strains. Full 16S rRNA genes of the most abundant sequences from each enrichment were blasted to NCBI database (Clark et al. 2016) for initial ID. Sequences with confirmed cyanobacterial identity, were further analyzed using the comprehensively curated Cydrasil (https://github.com/FGPLab/cydrasil) database of cyanobacterial 16S rRNA diversity by placement on a pre-calculated tree.

#### Data Summary

In keeping with FAIR principles (Findable, Accessible, Interoperable, Reusable data), all analyses presented in this paper can be reproduced and inspected with the associated GitHub repository https://nico-chung.github.io/loopseq/.

## Results

### LoopSeq long-read sequencing

The Zymo community genomic DNA (gDNA) quantities from the eight bacterial species were evenly distributed by mass, and the species within the standard vary widely in genome size, G+C content and 16S ribosomal copy number (GCN). Five of the eight Zymo strains were replaced by the manufacturer with similar strains, starting with Lot ZRC190633. For this study, we used a Zymo mock community sample after this update had been applied (Lot ZRC190811). While the Zymo mock community is a widely used standard in microbiome studies some of the references provided do not perfectly match the strains included in the standard (McGovern *et al*., 2018). As noted in other analyses of the Zymo standard, the complete sequences of all the 16S rRNA genes of each strain present in the mock community are not disclosed by the manufacturer and the reference sequences that are provided do not exactly correspond to the actual 16S rRNA gene sequences provided in the standard (Nicholls *et al*., 2019). Nevertheless, full-length 16S contigs are sufficiently discerning to allow for mapping to the correct species references. The whole genome references that contain all 16S rRNA gene copies in the genome were used to ensure greater mapping specificity, since individual 16S rRNA gene copies within a bacterial genome can diverge in their sequence (Sun *et al*., 2013).

The American Type Culture Collection (ATCC) offers extensive support materials that characterize their mock community samples (Biodefense and Emerging Infections Research Resource Repository (BEI Resources), VA, USA). Two mock community samples were sequenced here, the first contains 12 bacterial species endemic to human gut (‘ATCC-gut’; MSA-1006), and the second contains six bacterial species commonly found in the human oral cavity (‘ATCC-oral’; MSA-1004). Many of the species in the ATCC mock communities can be linked to exact strain references including assembly details and genome annotations. Whenever strain references were not available from the ATCC Genome Portal (https://genomes.atcc.org/), we used the strain name to find identical or similar strain references in other genome repositories (i.e. NCBI RefSeq CG database).

Both ATCC-gut and ATCC-oral (Table S1) mixes contain equimolar quantities of gDNA of their constituent species. The expected distribution of 16S rRNA genes per species in the community is calculated based on the number of 16S gene copies and the genome size of each species. For these mock communities, exact strain names were known. For all six strains in the ATCC-oral community whole genome references from the ATCC Genome Portal were used. For the 12 strains in the ATCC-gut community, whole genome references from the ATCC Genome Portal were used for six strains and exact strain references for the remaining species were obtained from NCBI RefSeq CG.

Extracted DNA samples were processed into sequence-ready libraries using LoopSeq library preparation kits that capture 16S ribosomal DNA (rDNA) targets (Loop Genomics, San Jose, CA, USA). The core principle behind LoopSeq technology involves attaching a unique molecular identifier (UMI) DNA tag to the end of each ‘parent’ molecule in a sample. Additionally, a sample-specific tag (i.e. Loop Sample Index) is incorporated into each molecule from the same physical sample. Through SLR sample prep chemistry, the UMI is distributed to random positions within each parent molecule but not to other molecules. After this step, samples are multiplexed into one reaction tube and molecules that contain a UMI tag are then broken into smaller units at the junction adjacent to the UMI, creating a library of UMI-tagged fragments that are compatible with a short-read Illumina sequencing. Short-reads that contain the same UMI were derived from the same parent molecule, and are binned and informatically reassembled back into the full-length sequence of the original molecule, producing a synthetic, high-quality, curated long-read that covers all 9 variable regions of the 16S rRNA gene.

Raw short-reads prepared with the LoopSeq kit were uploaded to the Loop Genomics cloud platform for processing (https://www.loopgenomics.com/16s-readcloud). Within this analysis pipeline, reads are first trimmed to remove adapter sequences that are not part of the original 16S molecule. The pooled reads are then de-multiplexed based on the Loop Sample Index attached to each read, which segregates reads based on the sample from which they originated, thus resolving the Loop multiplexing step. Next, sample-specific reads are grouped by UMI such that reads with an identical UMI are binned and reassembled with SPAdes genome assembler (Bankevich *et al*., 2012) to produce a synthetic long-read (i.e. a contig).

Each contig is assembled from a collection of short-reads with the same UMI, indicating their shared origin from a single parent molecule, with each short-read covering a different sequence region of the parent molecule. Given sufficient short-read coverage across the full length of a 16S rRNA gene molecule, it is possible to re-assemble the entire sequence of the original long molecule by linking reads. Short-reads whose sequences partially overlap can be linked through shared sequence identities, which can be arranged in the correct order for assembling the original long molecule sequence. Each assembled contig thus represents an original 16S rRNA gene. Contigs assembled with fewer reads results in shorter sequences with lower sequence accuracy. Assembled contigs that are full length are then mapped for classification against the SILVA database (Pruesse *et al*., 2007). The pipeline outputs both FASTA and FASTQ files that contain every contig generated regardless of mapping status, a CSV file that summarizes assembly statistics for each assembled contig by sample (including taxonomic classification), and a run summary in HTML format that provides charts for identified species per sample, species distributions, and other important statistics.

### LoopSeq long-reads accurately quantify a mock microbiome

After LoopSeq 16S sequencing of the Zymo mock community, a total of 38,629 unique molecular identifiers (UMIs) were assembled into 16S rRNA long-reads (mean length = 1,430 bp; median = 1,508 bp). We considered 27,123 of these (70.2%) to be meaningful long-reads after filtering for full-length 16S molecules (mean length = 1,474 bp; median = 1,472 bp). The long-reads were then mapped back to the references of the Zymo mock community which showed taxonomic ratios expected in the community (Table S2). All 27,123 full-length Zymo-Loop long contigs could be mapped to the NCBI RefSeq general bacterial database at approximately the expected species-level ratios (Table 4). Only 0.4% of the contigs (*n*=104 contigs) mapped to an incorrect species reference. Almost all these contigs mapped to the correct genus (*n*=103), with a single instance of mapping to an unexpected genus (*Klebsiella*). Similarly, as a positive control, Illumina short-read V3-V4 sequencing of the Zymo mock community produced 3,278,936 paired-end reads which were merged into 2,239,559 reads. Of these, 2,237,457 were mapped to the Zymo mock community at approximately expected ratios (Fig. 1 and Table S2).

**Table 1.** Class profiles of Zymo-Loop and Zymo-V3V4 sequencing.

| Class | Zymo-Loop | Zymo-V3V4 |
| --- | --- | --- |
| Match | 1.00E+00 | 9.93E-01 |
| Mismatch | 7.48E-05 | 7.49E-03 |
| Deletion | 2.39E-06 | 8.37E-05 |
| Insertion | 2.15E-06 | 3.12E-05 |

**Table 2.** Mapping ATCC-gut to exact strain whole genome references and classification of ATCC-gut contigs using Kraken 2.

| Species | Expected (%) | Strain | ATCC-gut (%) | Species-level classification? | Strain-level classification? | Strain detected |
| --- | --- | --- | --- | --- | --- | --- |
| *Bacteroides fragilis* | 8.0 | ATCC 25285 | 11.7 | Yes | No |  |
| *Bacteroides vulgatus* | 9.3 | ATCC 8482 | 10.7 | Yes | Yes | *Bacteroides vulgatus* ATCC 8482 |
| *Clostridioides difficile* | 16.0 | ATCC 9689 | 21.6 | Yes | Yes | *Clostridioides difficile* M68 |
| *Enterobacter cloacae* | 10.7 | ATCC 13047 | 7.6 | No | No |  |
| *Escherichia coli* | 9.3 | ATCC 700926 | 8.1 | Yes | Yes | *Escherichia coli* O7:K1 |
| *Salmonella enterica* subsp. *enterica* | 9.3 | ATCC 9150 | 4.3 | Yes | Yes | *Salmonella enterica* subsp. *Enterica* & *Salmonella enterica* subsp. *Salamae* |
| *Bifidobacterium adolescentis* | 6.7 | ATCC 15703 | 0.3 | Yes | Yes | *Bifidobacterium adolescentis* ATCC 15703 |
| *Enterococcus faecalis* | 5.3 | ATCC 700802 | 6.4 | Yes | No |  |
| *Lactobacillus plantarum* | 6.7 | ATCC BAA-793 | 6.1 | Yes | No |  |
| *Helicobacter pylori* | 2.7 | ATCC 700392 | 5.8 | Yes | Yes | *Helicobacter pylori* Shi169 |
| *Yersinia enterocolitica* | 9.3 | ATCC 27729 | 7.4 | Yes | No |  |
| *Fusobacterium nucleatum* subsp. *nucleatum* | 6.7 | ATCC 25586 | 9.8 | Yes | Yes | *Fusobacterium nucleatum* subsp. *nucleatum* ATCC 25586 |

**Table 3.** Mapping ATCC-oral to exact strain whole genome references and classification of ATCC-oral contigs using Kraken 2.

| Species | Expected (%) | Strain | ATCC-oral (%) | Species-level classification? | Strain-level classification? | Strain detected |
| --- | --- | --- | --- | --- | --- | --- |
| *Schaalia odontolytica* | 11.5 | ATCC 17982 | 3.9 | Yes | No |  |
| *Prevotella melaninogenica* | 15.4 | ATCC 25845 | 13.7 | Yes | No |  |
| *Fusobacterium nucleatum subsp. nucleatum* | 19.2 | ATCC 25586 | 28.1 | Yes | Yes | *Fusobacterium nucleatum subsp. nucleatum* ATCC 25586 |
| *Streptococcus mitis* | 15.4 | ATCC 49456 | 15.4 | Yes | No |  |
| *Veillonella parvula* | 15.4 | ATCC 17745 | 13.4 | Yes | No |  |
| *Haemophilus parainfluenzae* | 23.1 | ATCC 33392 | 25.6 | Yes | No |  |

**Table 4.** Species-level ratios after mapping Zymo-Loop to a general whole genome database containing 35,494 bacterial species.

| **Species** | **Expected relative abundance (%)** | **Zymo-LoopSeq 16S (%)** |
| --- | --- | --- |
| *Bacillus subtilis* | 17.4 | 21.0 |
| *Enterococcus faecalis* | 9.9 | 9.7 |
| *Escherichia coli* | 10.1 | 10.9 |
| *Lactobacillus fermentum* | 18.4 | 11.3 |
| *Listeria monocytogenes* | 14.1 | 17.4 |
| *Pseudomonas aeruginosa* | 4.2 | 4.2 |
| *Salmonella enterica* | 10.4 | 11.2 |
| *Staphylococcus aureus* | 15.5 | 14.0 |
| Other species | 0.0 | 0.4 |

**Figure 1.**
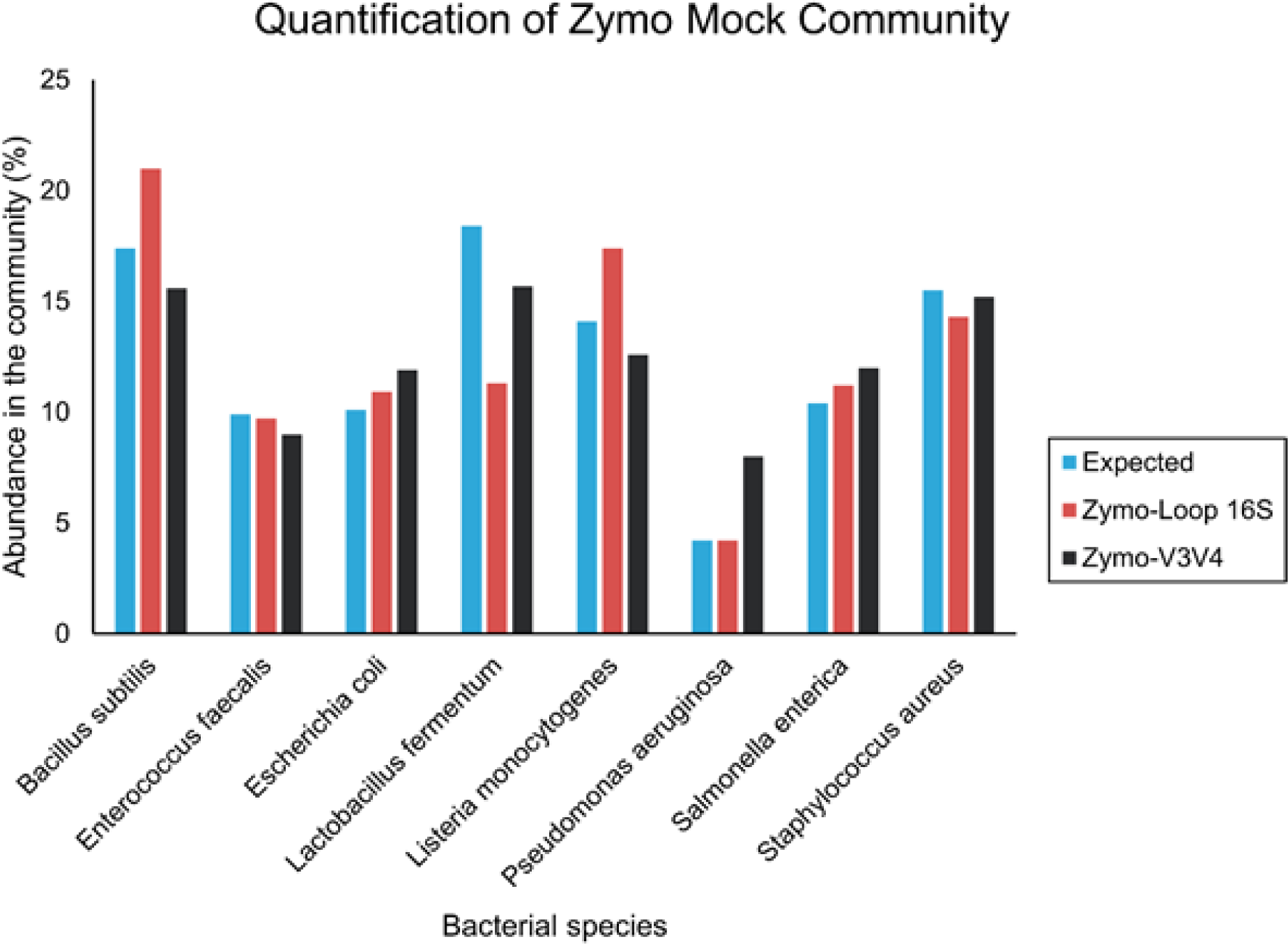
Quantification of Zymo mock community. Expected vs. observed abundance of known species for Zymo-Loop and Zymo-V3V4 sequencing inferred by mapping sequencing reads to known references provided by Zymo Research.

### LoopSeq long-reads identify a non-redundant 16S rRNA gene copy number ratio signature

LoopSeq can accurately resolve 16S rRNA GCNs from mixed microbial populations. As a proof of concept, we analyzed the GCNs within the *B. subtilis* genome from the Zymo community. A total of 5,692 Zymo-Loop full-length long-reads and 348,730 Zymo-V3V4 short-reads that initially mapped to the *B. subtilis* strain B-354 genome from the Zymo community were re-mapped to a database containing only non-redundant *B. subtilis* 16S rRNA gene sequences. *B. subtilis* B-354 contains the highest copy number of 16S rRNA genes (*n*=10, Table S2) per genome of any strain in the Zymo sample. Multiple sequence alignment consolidated the genomic 16S copies into 5 unique 16S rRNA gene sequences (full-length exact sequence variants or ESVs). The first group contains 5 members comprising 16S rRNA gene copies #1, #4, #8, #9 and #10 (which are identical), the second has two members (identical copies #2 and #3), while the remaining three copies (#5, #6, #7) are all unique to one another and to the other 16S rRNA groups (Fig. 2*a*). We used this information to examine if the intragenomic 16S rRNA gene copy ratios were accurately reported by LoopSeq SLR contigs. Correct mapping to these unique, non-redundant copies should produce a mapping ratio of 5:2:1:1:1. The Zymo-Loop long-read data was derived from experimental sampling of 10 ng of gDNA (which contains thousands of bacterial genomes) and reflected the actual ratio of non-redundant 16S rRNA gene copies contained within a single genome (5:2:1:1:1). In contrast, the results generated using Zymo-V3V4 short-reads yielded an erroneous 16S rRNA gene copy ratio (∼2:2:2:2:1) as the V3-V4 region does not capture many of the relevant base pair differences that exist among the 16S rRNA gene copies in this species.

**Figure 2.**
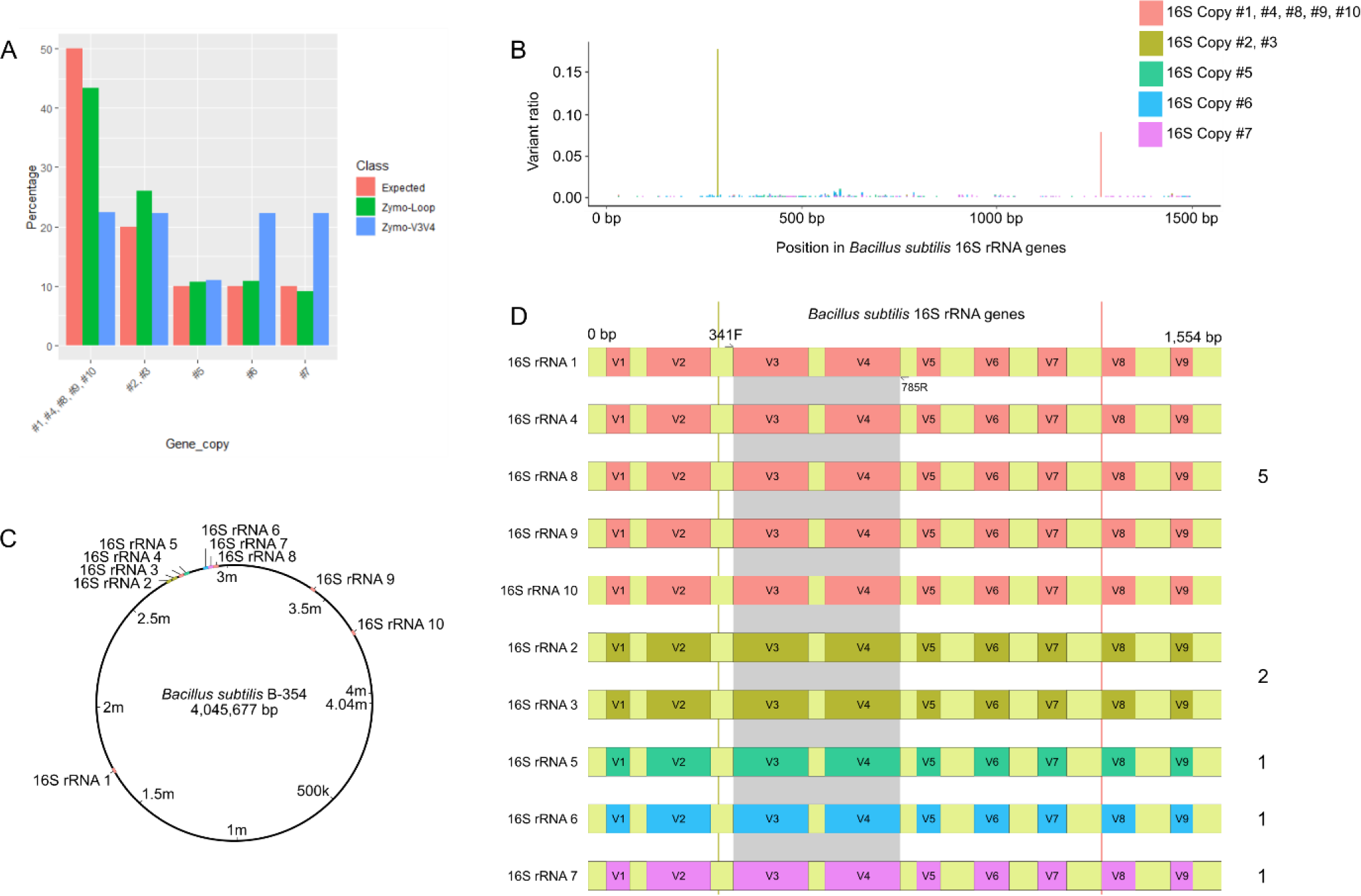
*Bacillus subtilis* gene copy numbers. **a**, Mapping Zymo-Loop, and Zymo-V3V4 *B. subtilis* sequences to non-redundant 16S gene references. b, Overlapping distribution of mismatch errors of Zymo-Loop contigs across all five non-redundant copies of the *B. subtilis* B-354 16S rRNA genes. Identified ESV are color-coded. **c**, Genomic placement of the 10 copies of the 16S rRNA gene within the *B. subtilis* B-354 genome, colored by ESV. **d**, Loop 16S detects 10 16S rRNA genes as 5 distinct full-length ESVs at a ratio of 5:2:1:1:1. Sequencing of the identical V3-V4 regions of the 16S rRNA gene in only sensitive enough to resolve a single ESV, and misses single base pair differences that distinguish the full-length ESVs.

### Extremely low error rates observed in LoopSeq SLR contigs

The *B. subtilis* B-354 genome contains 10 copies of the 16S rRNA gene of identical length (1,558 bp; Fig. 2*c*). Therefore, the full-length contigs generated from the Zymo-Loop sequencing data can be overlaid on the same plot to visualize base pair differences along their length (Fig. 2*b*). Most mismatch errors are randomly distributed along the length of the contig. The nucleotide differences that produced the two highest variant ratio peaks (gene copy #2 & #3 c. 285G>A and gene copy #1, #4, #8, #9 & #10 c.1268G>A) tentatively represent real variants in the non-redundant 16S rRNA genes as these exact same nucleotide mutations were previously identified (McIntyre *et al*., 2019). Consequently, the true mismatch error rate, after excluding these two false positive mutations which together account for 454 of 1691 total ‘mismatch’ errors, is very low (Fig. 2*d*). Altogether, the Zymo-Loop 16S dataset was at least an order of magnitude more accurate than Zymo-V3V4 across the frequency of mismatches, insertions, and deletions (Table 1).

### ATCC-gut sample: LoopSeq SLR results

After sequencing the ATCC-gut mock community, reads were assembled into 20,027 contigs (mean length = 1,376 bp; median = 1,462 bp). After filtering for full-length 16S rRNA genes, 9,087 contigs remained (45.4%) with a mean length of 1451 bp. Of these, 9,084 (99.9%) contigs could be mapped to one of the 12 reference sequences at approximately the expected ratios barring *Bifidobacterium adolescentis*, which was below expectation (Table 2). We speculate that this was the result of a sequencing artefact arising from an imperfect PCR primer match at the *B. adolescentis* primer region used to amplify the 16S rRNA gene (Frank *et al*., 2008). Using Kraken 2, all 9,087 contigs could be taxonomically classified (Table 2). Of the 12 species in the sample, 11 were classified correctly to the species level, of which seven were classified down to the strain level. Among the seven detected strains, three were exact matches to the ATCC strain designations.

### ATCC-oral sample: LoopSeq SLR results

After sequencing the ATCC-oral mock community, reads were assembled into 33,222 contigs (mean length = 1,372 bp; median 1,478 bp). After filtering for the full-length 16S rRNA gene, 16,392 contigs remained (49.3%) with a mean length of 1,460 bp and a median length of 1,462 bp. Kraken 2 taxonomically classified all 16,173 contigs (Table 3). Here, 16,172 contigs (99.9%) could be mapped to one of the six known references at approximately the expected ratios, although *Schaalia odontolytica* was substantially lower in proportion than expected (Table 3). Notably, all six species in the sample were detected and classified correctly to the species level. *Fusobacterium nucleatum subsp. nucleatum* could be further classified to the strain level (ATCC 25586) and was an exact match to the ATCC strain designation.

### Read depth required for contig assembly

The ability to reconstruct a long contig from overlapping short-reads requires, in addition to UMI tagging, sufficient read depth to cover the entire length of a 16S molecule with short-reads. Fig. 3 shows the relationship between the number of reads needed to generate a contig (of any length) and the fraction of total contigs (i.e. UMIs) that achieve full-length status. The x-axis marks individual UMIs that represent ∼9,000 original 16S molecules arranged from shortest to longest. The y-axis indicates the number of reads that shared the same UMI (binned by UMI) that were used to assemble that contig. The color indicates contig length (16S full-length is ∼1500 bp). Full-length contigs are displayed in yellow, clearly showing the reads/UMI transition point where the synthetic assembly achieves complete reconstruction of the 16S rRNA gene. This value is ∼100 reads/contig on average for LoopSeq 16S kits.

**Figure 3.**
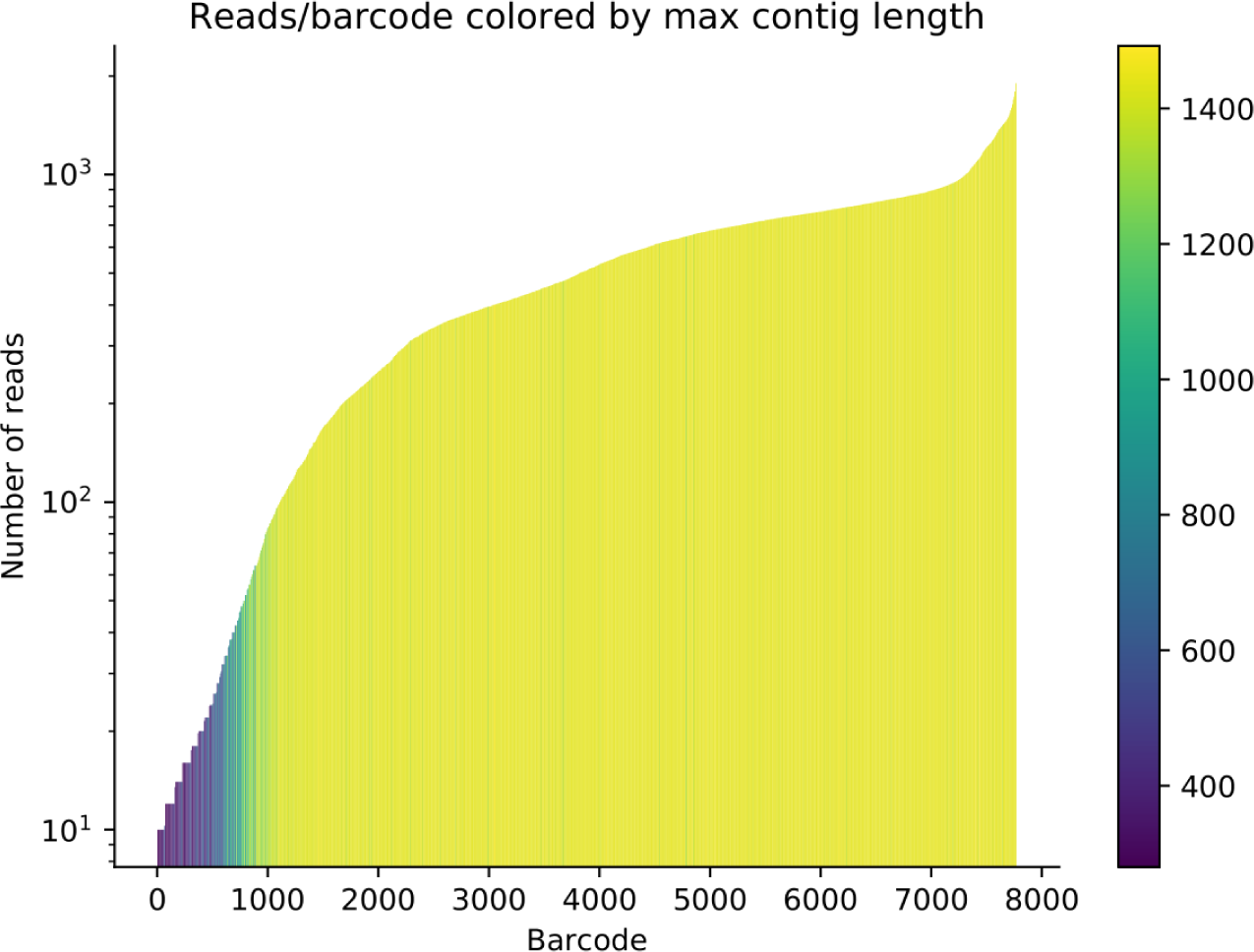
LoopSeq produces full-length contigs. Contig length histogram showing all LoopSeq contigs as a function of the number of short-reads required to assemble full-length contigs. The color legend indicates assembled contig length scaled from dark blue (short contigs) to bright yellow (long contigs).

### Water microbiome analysis

Water samples were collected from four different locations in Madison, WI, USA and processed using LoopSeq 16S kits. The resulting data sets were filtered to include only full-length contigs for the following samples: Aquarium (6,428 contigs), FeynmanPond (8,251 contigs), RainBarrel (4,673 contigs), and RivaPond (3,564 contigs). All LoopSeq SLR contigs across the four samples could be classified using Kraken 2 after running the standard database. Although many contigs were classified to the species-level, some contigs were classified at higher taxonomic levels. This did not permit a comparison of relative species abundance between samples. To obtain an approximation of species abundance, Bracken was used to redistribute contigs to the species level. The total species counts and top three ranking species by abundance for each sample are shown in Table 5 and a rank abundance plot in Fig. 4*a*. The Aquarium sample had the lowest species diversity and the most uneven species distribution as it was dominated by *Limnohabitans* sp. By contrast, the FeynmanPond sample was the most evenly distributed sample and had the highest species diversity (Fig. 4*b*).

**Table 5.**
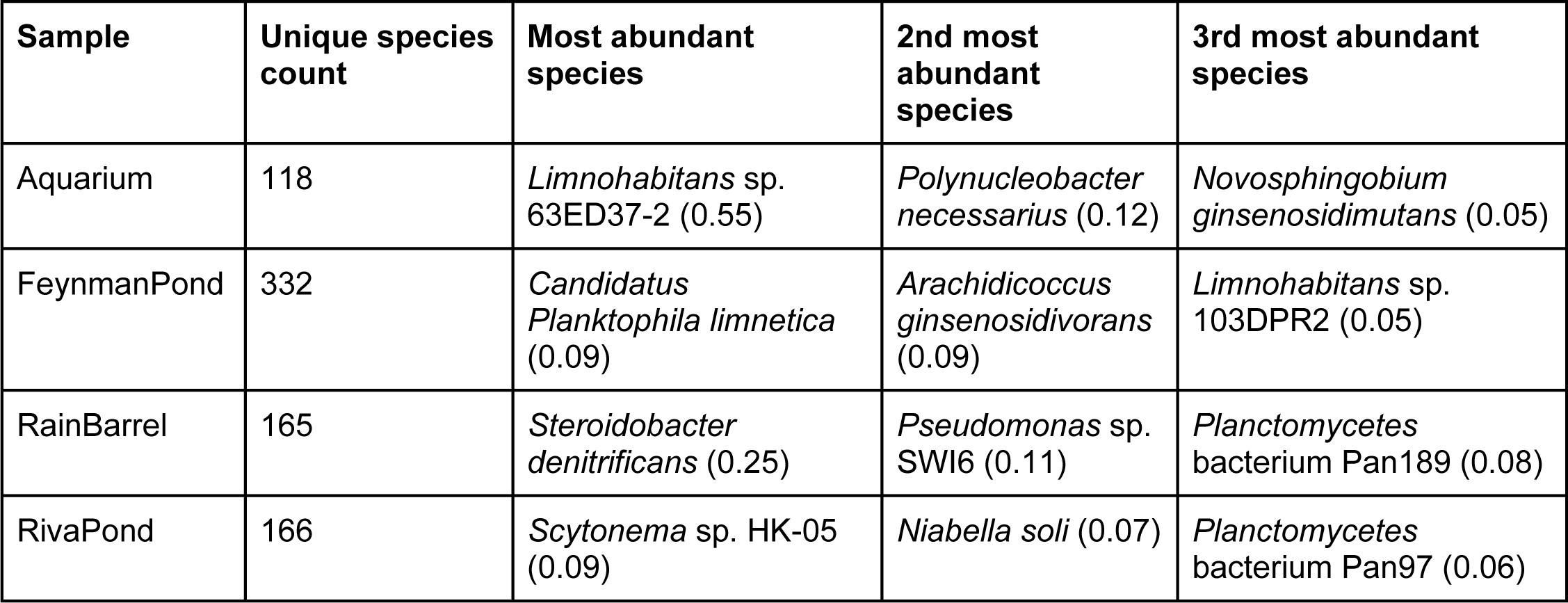
Water samples with species counts and top three ranked species by abundance, with the relative abundance for each species shown in brackets.

**Figure 4.**
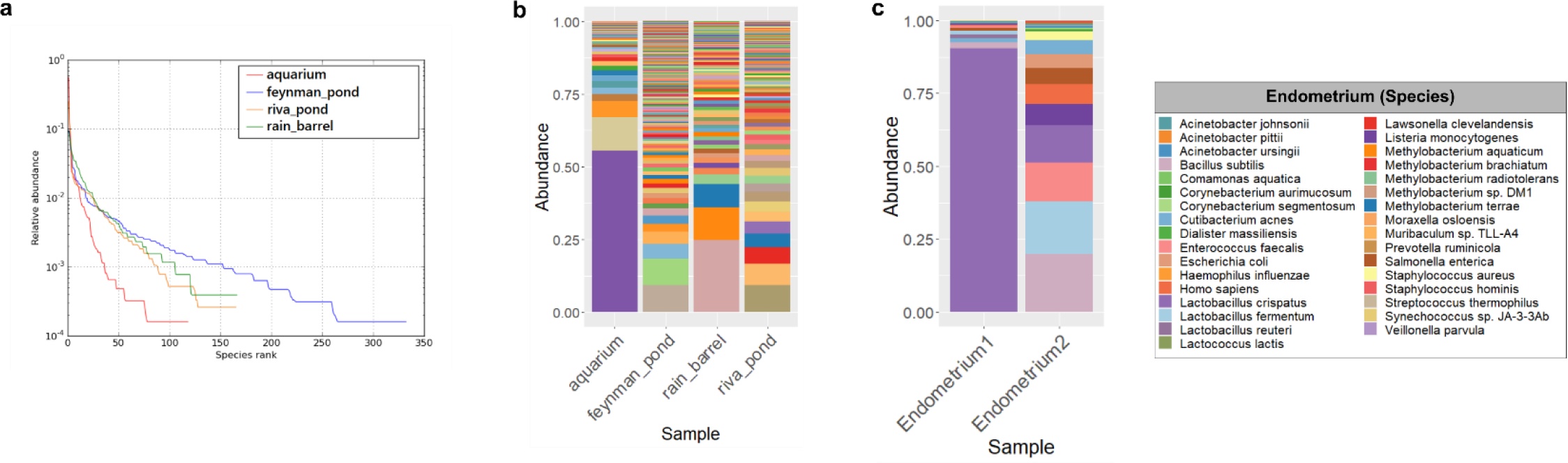
Microbiome analyses using 16S rRNA gene sequencing via SLR. **a**, Species rank abundance plot of water samples. The Aquarium sample had the lowest species diversity and the most uneven species distribution. The Feynman Pond sample was the most evenly distributed and had the highest species diversity. **b**, Bar chart of species-level relative abundance for each aquatic sample **c**, Relative abundances of bacterial species within the endometrium of two patients exhibiting different clinical presentations relating to complications with conception.

### Human uterine lavage microbiome analysis

With the use of an optimized method for DNA extraction (see supplemental methods), uterine lavage samples were collected from two patients exhibiting complications with conception. Samples were prepared for LoopSeq 16S processing to investigate the bacterial composition of the dysbiotic endometrium. Previous work with mock communities validated the efficacy and reliability of this method (see Results above and Supplemental Results). Thus, we isolated patient samples and processed them using the LoopSeq workflow and Loop Genomics cloud platform-powered taxonomic analysis.

Patient 1 exhibited secondary sterility (3 spontaneous pregnancy losses and 6 IVF attempts) which may manifest in the presence of a dysbiotic microbiome composition (Franasiak *et al*., 2016). The uterine lavage community was dominated by a single *Firmicutes* species. More than 90% of full-length 16S rRNA genes were assigned to *Lactobacillus crispatus* (Fig. 4*c*, endometrium1). Patient 2 also had a history of repeated, unsuccessful attempts at conception, yet presents a very different microbiome composition: 19.8% *Bacillus subtilis*, 18.2% *L. fermentum*, 13.2% *Enterococcus faecalis*, 12.7% *L. crispatus*, 7.4% *Listeria monocytogenes* and 5.7% *Salmonella enterica* (Fig. 4*c*, endometrium2).

### Cyanobacterial cultures from biological soil crust

For the two cyanobacterial enrichments, we targeted the establishment of uni-cyanobacterial cultures from motile filamentous cyanobacteria (i.e. *Microcoleus vaginatus* and those in the *Microcoleus steenstrupii* complex) that pioneer the formation of biocrust communities in arid lands, particularly in the Southwestern US. Initial identification of the cyanobacteria present in each enrichment was performed by means of Sanger sequencing of the 16S rRNA gene using cyanobacteria-specific primers (CYA359F/CYA781R; (Nübel *et al*., 1997). This yielded (∼600 bp) single sequences for each of the two enrichments: HSN023 (sample 7) and CYAN3 (sample 16). Both were then classified as belonging to the *M. steenstrupii* complex of genera. After LoopSeq sequencing however, we uncovered that HSN023 was a) not uni-cyanobacterial and b) contained significant amounts of heterotrophic contaminants. Of the 1703 contigs obtained, 69% were cyanobacterial, the remaining 31% assigned to non-cyanobacterial contaminants. We could resolve three cyanobacterial ESVs, assignable to two distinct family-level clades (Fig. 5).

**Figure 5.**
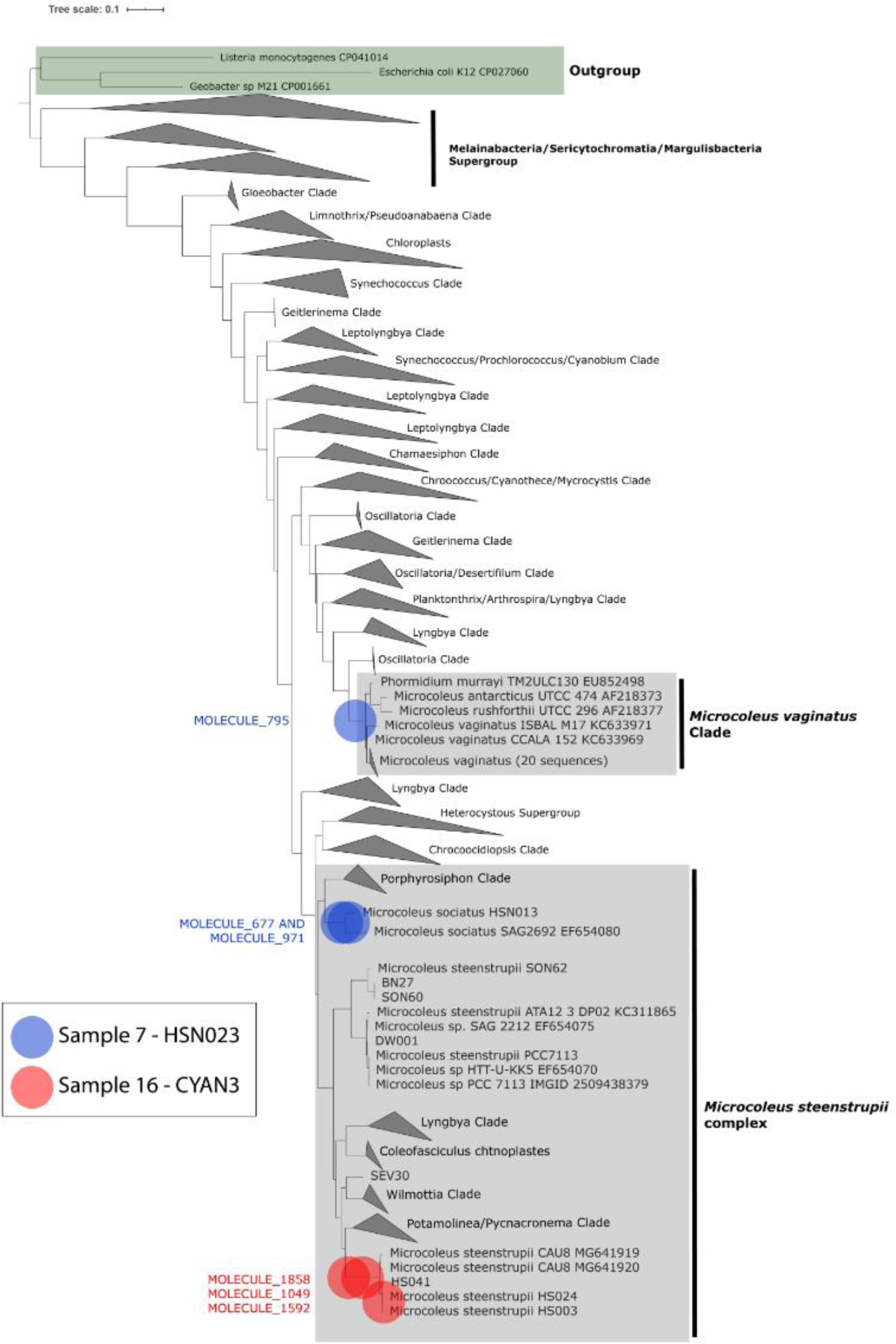
Phylogenetic relationships of biocrust isolates are accurately resolved. Biocrust Sample 7 (HSN023) and Sample 16 (CYAN3) cyanobacterial full-length 16S rRNA ESVs in resolved into 3 distinct cyanobacterial ASVs.

Two of the ESV’s, which accounted for the majority of contigs, were assignable to the *Microcoleus sociatus* clade (a member of the *M. steenstrupii* complex; Fernandes *et al*., in prep) and the third, which accounted for some 18% of cyanobacterial contigs, to the *M. vaginatus* clade (Fig. 5). *Post-hoc* microscopic inspection of HSN023 confirmed the presence of low-level populations of *M. vaginatus* morphotypes. The two *M. sociatus* ESVs (molecules 677 and 971) were very similar to each other (98.7%) and occurred at a frequency ratio very close to 1:1 among contigs, suggesting they represent variants of multiple ribosomal operons within a single genome. Hence HSN023 was composed of two cyanobacterial strains, with one having two slightly divergent ribosomal operons. In the case of enrichment (CYAN3 - sample 16) LoopSeq resolved two ESV’s accounting for 67% of all contigs. Two were 99.73 % similar (one a bit shorter) and we take it to be different molecules of the same sequence, and their sum was close to a 1:1 (56/45) proportion to the third ASV, likely representing two distinct operons within a genome. We interpret these results as confirming the uni-cyanobacterial status of the enrichment.

## Discussion

### LoopSeq SLRs have improved mapping and reduced errors

Previous work has shown that substantial improvements in species and strain-level taxonomic resolution are achieved by analyzing the full-length 16S rRNA gene compared to ∼500 bp regions of the gene (Martijn *et al*., 2019) and that ultra-low error rates are required in order to cluster long-read microbiome sequencing data into ESV’s (Callahan *et al*., 2019). LoopSeq technology uses unique molecular identifiers (UMIs) for each parent 16S rRNA molecule, which allows accurate long-reads to be constructed from amplicon sequencing data, overcoming the critical constraint of amplicon sequencing, i.e. short read lengths, by sequencing the entire 1.5 kb 16S rRNA gene and providing at least an order-of-magnitude lower error rates compared to existing short and long read modalities in reading microbial community DNA.

### Identifying intragenomic 16S variation using LoopSeq SLR contigs

The initial mapping of Zymo-Loop and Zymo-V3V4 to a known sample database quantification differences from the expected abundances for both approaches (Fig. 1). However, where the two methodologies differ greatly is in their resolution of intragenomic variation. We observed the most striking example among the 10 *B. subtilis* B-354 16S rRNA gene copies, where many single nucleotide differences were detected along the length of the 16S rRNA gene, and critically, not within the V3-V4 region of the gene (range from positions 348-792). Within the V3-V4 region, only copy #5 is unique (pos: 373 C>A) from copies #1, #2, #6 and #7, which are identical. As such, reads corresponding to these identical sequences map equally well to any of the 16S gene copies, and thus can be randomly assigned to a reference. This results in incorrectly assigning equal ratios for copies #1, #2, #6 and #7 after V3-V4 sequencing, in contrast to the correct 5:2:1 ratio assignment using the Loop-Zymo long-read data (Fig. 2a).

These results highlight a major shortcoming in sequencing hypervariable regions with short-reads alone; they do not permit an accurate assessment of intergenomic 16S rRNA gene ratios as the necessary sequence information required to distinguish unique copies from one another is missing. This limits the use of short-read sequencing for inferring bacterial subtypes based on their intergenomic 16S gene copy ratios. In general, the use of short hypervariable regions (V1V2, V1-V3, V4, etc.) has limited utility in identifying prokaryotes to the species leve<u>l</u> (Johnson *et al*., 2019) which is consistent with the results presented here.

### Simplified error reduction: correction by consensus

Another major advantage of the LoopSeq technology for accurate 16S rRNA gene sequencing is the ability to correct for sequencing errors at nucleotide resolution through base-position consensus. As shown in Table 1, the error rates for Loop-Zymo data were extremely low considering these long contigs have only been filtered to be full-length 16S rRNA genes, with no additional error-reducing methods applied. By all metrics, including rates of indels and mutations, the SLR contigs are at least an order-of-magnitude more accurate than traditional short-read sequencing alone. Even though SLR contigs were not quality filtered, denoised or processed using algorithmic error correction, the indel rates were extremely low, thereby increasing the confidence in classifications at the species-or strain-level.

These extremely low error rates stem from correction by consensus. At any nucleotide position where multiple short-reads overlap, a majority base call from all the short reads that span that position is used to arrive at the most common call, which filters out non-systematic errors (e.g. polymerase errors during PCR). Correction by consensus allows for a more faithful identification of which microbes are in a population by reducing false-positives, such that genuine sequencing errors are not counted as unique 16S molecules, which would artificially inflate the number of predicted species. Lower error rates are particularly important for reliably detecting 16S sequences of low abundance, which are hard to distinguish from sequencing errors/artifacts. The number of UMIs per sample does not impact the fidelity of a consensus call, since UMI-tagged fragments are assembled independently from different clusters of overlapping short reads. Lastly, since consensus accuracy is contingent on read depth, higher per base accuracy and a higher proportion of full-length 16S contigs is expected with increased sequencing yield.

Based on our comparative analysis between full-length SLR and V3V4 PE300 Illumina sequencing of the 16S rRNA gene we found that 16S SLRs provide: 1) significantly reduced error rates, which affect false positive identification rates, 2) accurate quantification of bacterial communities and 3) the ability to detect intragenomic unique 16S gene copy ratios. Furthermore, we found that despite the denoising of short read sequencing using e.g. DADA2 (Callahan *et al*., 2016), UNOISE2 (Edgar, 2016) and Deblur (Amir *et al*., 2017), as well as clustering similar sequences into OTUs and using *k*-mer based LCA methods, short read sequences do not contain sufficient information to consistently and accurately resolve microbiome communities with species or strain-level resolution in most cases. Our analysis shows that directly mapping full length LoopSeq SLRs that span all nine hypervariable regions to a reference database without any denoising it is possible to classify bacteria within a complex sample down to the species level with high accuracy.

While the application of denoising techniques to LoopSeq SLRs could potentially improve on these results even further our analysis has shown that even in the absence of such methods, analyzing full-length 16S SLRs is sufficient for precise classification. Bacterial genomes harbor many 16S rRNA gene copies that differ at specific nucleotide positions across multiple hypervariable regions, spanning distances much longer than the length of short read sequencing. Long reads that span all 9 hypervariable regions in a single molecule readout can detect variation along the entire set of 9 variable regions and capture the intragenomic species/strain signatures encoded in 16S gene ratios (Fig. 3*C*). Another critical of NGS methods is the inadvertent introduction of bias during sample preparation. Methods that rely on PCR amplification, which includes LoopSeq as well as short-read V3V4 prep kits, introduce potential biases as certain molecules can amplify with variable efficiently compared to others. NGS methods that employ UMI’s to do so by back-tracking to the starting 16S molecule counts at UMI tagging. LoopSeq SLR technology employs UMIs and negates at least some of this amplification bias. Post-sequencing short read data is collapsed such that short reads that share a UMI (once binned) represent the sequence of a single long read. Bias negatively impacts the detection of low abundance species (i.e. those constituting <1% of the community), including those found in complex and highly diverse environmental samples shown in this study (e.g. FeymanPond sample; Fig. 3A).

### Characterization of complex systems benefits from LoopSeq SLRs

#### Water samples

Aquatic environments are teeming with complex, dynamic microbial communities that influence global carbon cycling by absorbing ∼30% of anthropogenically produced carbon dioxide (Kottmeier *et al*., 2016). Due to their high phylogenetic complexity (Fig. 3*B*), analyzing water samples served as a good testbed for verifying the limits of accurate detection using LoopSeq 16S sequencing. The aquarium sample was dominated by a single species that accounted for ∼50% of reads, yet the entire community was much more complex than the simplified mock communities. The most diverse sample, FeymanPond, contained hundreds of unique species, of which the majority were considered rare at ≤ 2% relative abundance in the population. LoopSeq SLRs were able to faithfully reproduce the nuances of a complex microbial community while providing accurate classifications of microbes to the species or even strain level throughout the gradient of community complexity.

#### Uterine lavage microbiomes

The human endometrium microbiome has been the focus of several recent clinical studies, with goals of elucidating the role of microbiome composition in reproduction (Bracewell-Milnes *et al*., 2018). Notably, srNGS data from patients having undergone *in vitro* fertilization (IVF) suggest that the microbial composition of the endometrium can serve as a biomarker of reproductive health. Specifically, the phylum *Firmicutes* is frequently the most abundant in the endometrium, with *Lactobacillus* the dominant genus in the healthy uterus (Moreno *et al*., 2016, D’Ippolito *et al*., 2018). In pathological states, such as recurrent pregnancy losses, chronic endometritis, or in recurrent embryo implantation failure, there is a significantly reduced representation of *Lactobacillus* with a concomitant increased prevalence of pathogenic genera such as *Streptococcus, Staphylococcus*, and the family *Enterobacteriaceae* (*Gammaproteobacteria*) (Franasiak *et al*., 2016, Franasiak & Scott, 2017).

Our data corroborate previous research on the microbial communities in the human vagina (Brooks *et al*., 2017) indicating that remarkable differences in community can manifest in the uterine lavage of two patients with a history of complications with conception. Previous work has shown that *Lactobacillus crispatus* can dominate in these niches under healthy conditions (Ravel *et al*., 2011) and our work further implicates *L. crispatus* and *L. fermentum* as prominent colonists following progression to a dysbiotic state. Accurately resolving species and strains within the endometrium with the use of longer and more accurate sequencing reads may inform strategies for combating dysbiosis. As we move beyond basic community characterizations to linking bacterial populations to human disease or dysbiosis, the next generation of sequencing technologies will need to expertly characterize the species (or ideally strains) that become prevalent alongside a clinical indication. From the studies conducted here, the LoopSeq method delivered species-level resolution of the bacterial populations in clinical uterine lavage samples.

#### Analyses of enrichment cultures from biocrusts

Biological soil crusts are organosedimentary microbial assemblages that colonize the soil surface of arid lands (Belnap *et al*., 2016). These photosynthetically driven microbial communities physically stabilize the soil surface and thus provide a habitable environment for further microbial colonization (Garcia-Pichel & Wojciechowski, 2009, Couradeau *et al*., 2019) increasing carbon and nitrogen cycling (Elbert *et al*., 2012) in nutrient-poor arid lands soils. Filamentous cyanobacterial such as *Microcoleus vaginatus* and those in the *M. steenstrupii complex* are pioneer species that initiate biocrust formation across the continental US. Their targeted isolation in uni-cyanobacterial cultures is a prerequisite to produce inoculum in efforts to restore biocrust communities (Giraldo-Silva *et al*., 2019, Roncero-Ramos *et al*., 2019, Giraldo-Silva *et al*., 2020) under ecological pressure from human activities (Zaady *et al*., 2016). Because biocrust cyanobacterial components are biogeographically distinct (Garcia-Pichel *et al*., 2013) and are adapted to local conditions (Muñoz-Martín *et al*., 2019, Giraldo-Silva *et al*., 2020), inoculation must be done using local strains (Ayuso *et al*., 2017, Giraldo-Silva *et al*., 2019) to increase success rates (Giraldo-Silva *et al*., 2020). In this study we showed that LoopSeq SLR’s enabled the characterization of potential sources of inoculum with respect to identity and purity of cultured materials. For example, highly resolved identification of different 16S ASV’s turned out to be critical in detecting the polyclonal nature of the cyanobacterial enrichment HSN023, which lead to its exclusion from downstream use and its targeting for further purification. Studies utilizing the identification of full-length ASV’s and 16S gene copy number variation will be increasingly used as an additional layer of information for strain detection based on gene identity and copy ratios, as demonstrated for HSN023 and also potentially for CYAN3.

## Conclusion

In both mock communities and environmental datasets, LoopSeq 16S sequencing presents improved performance for obtaining accurate, full-length 16S rRNA genes from complex microbiomes. Our results show that synthetic long-read sequencing (SLR) outperforms short-read sequencing at documenting taxonomic diversity based on the 16S rRNA gene, both in synthetic and real-world data. In the aquatic experiment, SLR resolved taxonomy to the species level (if not the strain level) and was effective across a gradient of community complexities. In the uterine lavage study, SLR accurately identified indicator strains affiliated with dysbiosis following repeated misconceptions. We show in the example with cyanobacterial cultures from biocrust communities that SLR can reliably describe the strain (s) composition of bacterial enrichments, as well as to reveal the intra-genomic distribution of their 16S rRNA gene copies. Furthermore, SLR had extremely high accuracy across the 16S rRNA gene and ultimately allowed genuine transition mutations to be detected among the 10 16S rRNA gene copies within the *B. subtilis* genome. Finally, SLR faithfully reproduced the expected copy ratio of *B. subtilis* 16S genes while read length limitations of short-read sequencing failed to capture the segments of the 16S gene that were critical to capture the identity and copy number ratios of the different 16S genes within a genome. Collectively, our results indicate that synthetic long-reads generate sequencing reads that have the potential to lead to a more accurate and finely-resolved microbiome community characterization.

