## Supplementary Materials for "Accurate Microbiome Sequencing with Synthetic Long Read Sequencing"

**List of Tables**

**Supplementary Table 1.** Bacterial species in the ATCC-gut and ATCC-oral mock community samples.

**Supplementary Table 2.** Mapping Zymo-Loop and Zymo-V3V4 to references provided by Zymo Research.

**Supplementary Table 1.** Bacterial species in the ATCC-gut and ATCC-oral mock community samples.

| **Species** | **Strain** | **System** | **Genome Size (Mb)** | **16S rRNA gene copies** | **Expected proportion (%)** |
|---|---|---|---|---|---|
| *Bacteroides fragilis* | ATCC 25285 | Gut | 5.2 | 6 | 8.0 |
| *Bacteroides vulgatus* | ATCC 8482 | Gut | 5.2 | 7 | 9.3 |
| *Clostridioides difficile* | ATCC 9689 | Gut | 4.3 | 12 | 16.0 |
| *Enterobacter cloacae* | ATCC 13047 | Gut | 5.6 | 8 | 10.7 |
| *Escherichia coli* | ATCC 700926 | Gut | 4.6 | 7 | 9.3 |
| *Salmonella enterica subsp. enterica* | ATCC 9150 | Gut | 2.7 | 7 | 9.3 |
| *Bifidobacterium adolescentis* | ATCC 15703 | Gut | 2.0 | 5 | 6.7 |
| *Enterococcus faecalis* | ATCC 700802 | Gut | 3.2 | 4 | 5.3 |
| *Lactobacillus plantarum* | ATCC BAA-793 | Gut | 3.3 | 5 | 6.7 |
| *Helicobacter pylori* | ATCC 700392 | Gut | 1.7 | 2 | 2.7 |
| *Yersinia enterocolitica* | ATCC 27729 | Gut | 4.6 | 7 | 9.3 |
| *Fusobacterium nucleatum subsp. nucleatum* | ATCC 25586 | Gut | 2.2 | 5 | 6.7 |
| *Schaalia odontolytica* | ATCC 17982 | Oral | 2.4 | 3 | 11.5 |
| *Prevotella melaninogenica* | ATCC 25845 | Oral | 1.8 | 4 | 15.4 |
| *Fusobacterium nucleatum subsp. nucleatum* | ATCC 25586 | Oral | 2.2 | 5 | 19.2 |
| *Streptococcus mitis* | ATCC 49456 | Oral | 1.9 | 4 | 15.4 |
| *Veillonella parvula* | ATCC 17745 | Oral | 2.2 | 4 | 15.4 |
| *Haemophilus parainfluenzae* | ATCC 33392 | Oral | 2.1 | 6 | 23.1 |

**Supplementary Table 2.** Mapping Zymo-Loop and Zymo-V3V4 to references provided by Zymo Research.

| Species | Genome size (Mb) | GC content (%) | 16S rRNA gene copies | Gram stain | Expected proportion (%) | Zymo-Loop 16S (%) | Zymo-V3V4 (%) |
|---|---|---|---|---|---|---|---|
| *Bacillus subtilis* | 4.0 | 43.9 | 10 | + | 17.4 | 21.0 | 15.6 |
| *Enterococcus faecalis* | 2.8 | 37.5 | 4 | + | 9.9 | 9.7 | 9.0 |
| *Escherichia coli* | 4.9 | 46.7 | 7 | - | 10.1 | 10.9 | 11.9 |
| *Lactobacillus fermentum* | 1.9 | 52.4 | 5 | + | 18.4 | 11.3 | 15.7 |
| *Listeria monocytogenes* | 3.0 | 38.0 | 6 | + | 14.1 | 17.4 | 12.6 |
| *Pseudomonas aeruginosa* | 6.8 | 66.2 | 4 | - | 4.2 | 4.2 | 8.0 |
| *Salmonella enterica* | 4.8 | 52.2 | 7 | - | 10.4 | 11.2 | 12.0 |
| *Staphylococcus aureus* | 2.7 | 32.9 | 6 | + | 15.5 | 14.3 | 15.2 |
